# Aggression: A gut reaction? The effects of the gut microbiome on aggression

**DOI:** 10.1101/2023.10.26.564110

**Authors:** Atara Uzan-Yulzari, Sondra Turjeman, Dmitriy Getselter, Samuli Rautava, Erika Isolauri, Soliman Khatib, Evan Elliott, Omry Koren

## Abstract

Recent research has unveiled conflicting evidence regarding the link between aggression and the gut microbiome. In our investigation, we meticulously examined the behavioral patterns of four groups of mice – wild-type, germ-free (GF), mice treated with antibiotics, and recolonized GF mice – to gain mechanistic insights into the impact of the gut microbiome on aggression. We discovered a significant correlation between diminished microbiome and increased aggression. Importantly, this behavioral shift could be restored when a WT microbiota was reinstated. Microbiota manipulation also significantly altered brain function, particularly in aggression-associated genes, and urine metabolite profiles. Notably, our study extends beyond the murine model, shedding light on clinical implications of early-life antibiotic exposure. We found that fecal microbiome transplants from 1mo old infants prescribed antibiotics during their first days of life led to a marked increase in aggression in recipient mice. This research demonstrates that the microbiota modulates aggression and underscores its importance in the realm of behavioral science.

**One-Sentence Summary:** The antibiotic-altered gut microbiome is implicated in increased aggression. It also leads to altered brain function, particularly in genes linked to aggression, and urine metabolite profiles showing a multi-system response to microbiota disruption.

---

Aggression is an intentional, complex, multifaceted social behavior, one prevalent in almost all species and intimately associated with host survival. Animals display aggression when defending territory, securing and defending food and mates, and establishing dominance hierarchies. The behavior is modulated by numerous factors including specific genes, neurotransmitters, environmental factors, pheromones and hormones(*1-4*), which are generally thought to be conserved across species ranging from fruit flies, to mice, to humans(*5, 6*). In mammals, where the aggression mechanisms appear to be more complex, it is thought that hormones play a significant regulatory role(*7*), and there is also emerging evidence that the gut microbiome may be involved.

Numerous studies have provided evidence supporting the notion of a gut-brain axis, with crosstalk mediated by the microbiome(*6*). While research on the role of the microbiome in aggression is in its infancy, connections have been identified in *Drosophila melanogaster*(*8*), dogs(*9*), mice(*10, 11*), and humans(*12*). The impact of changes in microbial composition on aggression is still unclear though, as one study found that an absence of bacteria led to decreased aggression in fruit flies(*13*). Similar findings have been made in hamsters(*14*), but several studies revealed increased aggression following antibiotic treatment or in germ-free settings(*8, 10, 11*). Many of these published studies are preliminary or descriptive, and to date, none have uncovered mechanisms or even explained the network of interactions between aggression, hormone levels, gene expression, metabolite profiles, and gut microbiome. Here we examine this complex network in a model organism using germ-free, antibiotic-manipulated, conventionalized, and humanized animals to reveal the importance of an intact microbiome in mediating host aggression.

## The role of the microbiome in mediating aggression

As recent research is inconclusive, we first aimed to examine the impact of gut bacterial population on aggression in a murine model. To this end, we compared behavior profiles (resident intruder test(*13*)) of germ-free (GF) mice, specific pathogen-free (SPF, wild type baseline) mice, antibiotic-treated SPF mice (ABX), and GF mice that were re-conventionalized with SPF fecal samples at age 5 weeks (C-GF). Three behavior assays were performed, each trial lasting 10 minutes (Figure 1A).

**Fig 1.**
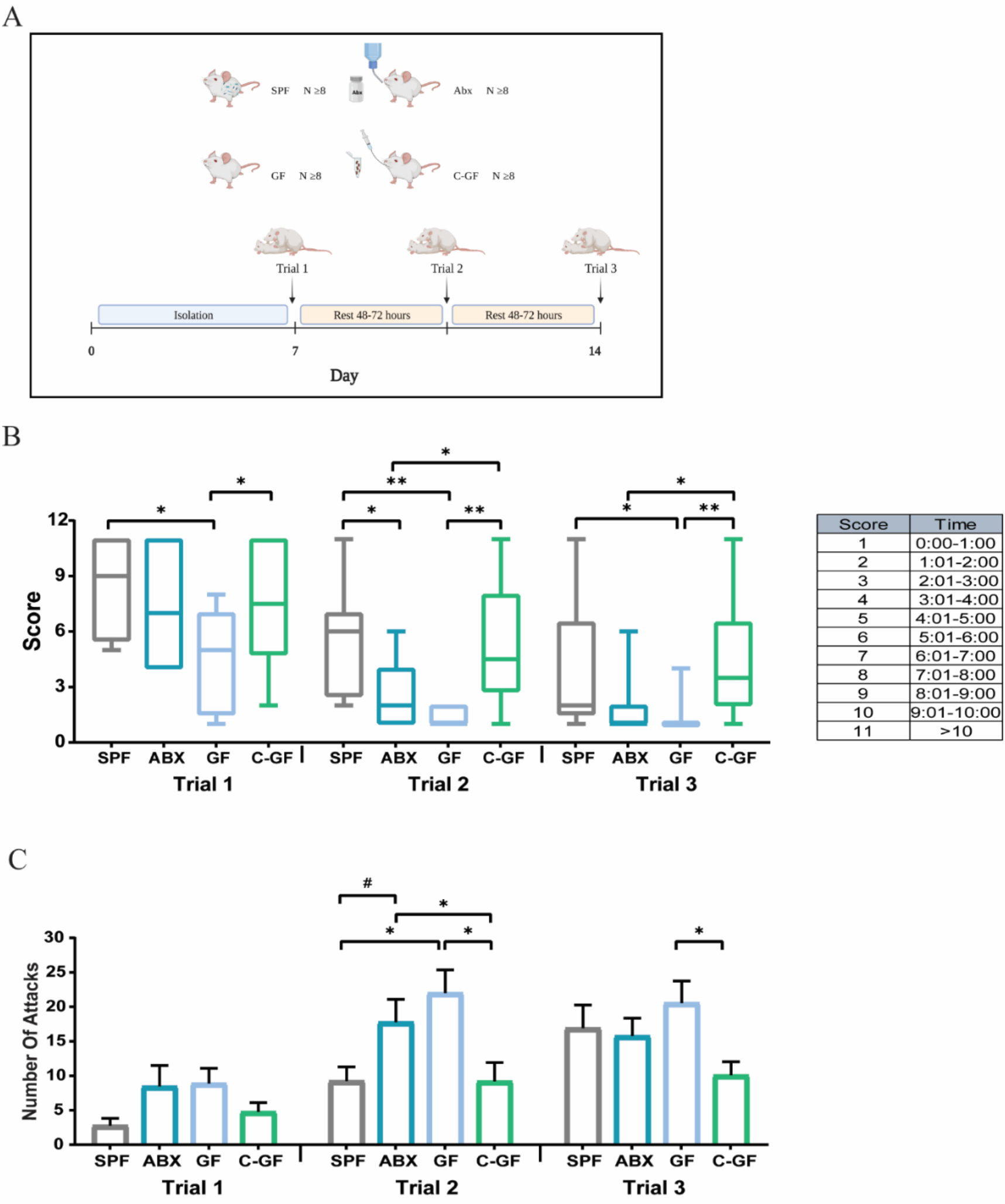
Gut microbiome modulates aggression in mice. (A) Experimental design – aggression was examined using the resident-intruder test. A resident male mouse was isolated with a female for 7 days and tested against an intruder male mouse three times at 2-4 days intervals. The female was removed 1 hour before the experiment, and each experiment lasted for 10 minutes. At the end of the experiment, the intruder was removed, and the female was returned to the cage. Aggression was measured using two parameters: (B) attack latency, the time between the first introduction of the intruder until the first attack by the resident, every minute received a score in order to achieve a discrete and non-continuous variable, and (C) number of attacks, the overall number of attacks in each trial (n≥9, ^#^*P*=0.06, ^*^*P*<0.05, ^**^*P*<0.01, number of attacks represents the mean +/- SEM).

The GF group demonstrated significantly higher aggression levels according to two parameters: they were faster to attack and attacked more frequently compared to SPF mice throughout the three trials (Fig 1B and 1C respectively). Moreover, the mean number of attacks among GF mice during the 10-minute trials was higher in all three trials compared to SPF mice, with statistically significant differences in the second trial (Fig. 1C). To investigate whether these differences resulted from the absence of bacteria, we manipulated the gut bacterial population by administration of antibiotic treatment to SPF mice, and in a similar manner, we colonized GF mice with stool samples from age-matched SPF mice. Consistent with the GF results, antibiotic administration led to elevation in aggression; latency to attack in antibiotic-treated mice was significantly lower compared to SPF in the second trial and similar to the GF group (Fig 1B). In addition, the number of attacks increased following antibiotic treatment (Fig. 1C). Interestingly, colonization of GF mice showed a significant reduction in aggression levels; C-GF mice were slower to attack (Fig 1B), and the number of attacks was significantly lower compared to GF groups (Fig. 1C).

## Mechanisms underlying increased aggression in ABX-treated mice

After revealing that absence of a microbiome (GF) or significant perturbations to the microbiome (ABX) increases aggression levels, we explored the web of interactions underlying this relationship. The primary mode of communication among mice is through scents, and urine serves as a main source of volatile and nonvolatile compounds. Analyzing urine composition can help identify potential compounds that might be linked to aggression, and thus we first examined the metabolite profile of mouse urine using untargeted metabolomics. Principal component analysis (PCA) was performed to evaluate the overall differences in the metabolic profiles between all groups prior to the aggression assay and between samples taken from SPF mice following the resident-intruder test (Fig 2). A clear separation can be seen between the SPF-T0 and the GF groups, based on the PCA plot. Moreover, antibiotic treatment of SPF mice led to a dramatic shift in the metabolic profile compared to the SPF-T0 group. Similarly, the metabolite profile of the C-GF mice shifted and was more similar to the SPF group rather than the GF group. When we examined all the groups together, there was no clear separation between the two time points of the SPF group: T0 and T14 (Fig 2A). However, when we performed PCA for only these two groups, we saw separation which was likely caused by the aggression trials (Fig 2B). Altogether, 99 metabolites from the urine samples (identified or putatively annotated) were differentially abundant (FDR *p*<0.05) in at least one of the statistical comparisons performed between the four experimental groups or nominally different (p<0.05) between T0 and T14 in the SPF group. Interestingly, we identified several metabolites that changed after antibiotics treatment and found they also changed after aggression trials in the same manner (Figure 2C-2H). Tryptophan (Trp), an essential amino acid that functions as a precursor of the neurotransmitter serotonin, and creatinine are examples of metabolites that change due to the absence or presence of a bacterial population; the GF group showed significantly higher levels of Trp and creatinine compared to the SPF group. Likewise, antibiotic administration to SPF mice, for a period of 2 weeks, led to elevated Trp and creatinine levels compared to the SPF group, whereas bacterial colonization of GF mice led to a reduction in Trp and creatinine levels (Fig. 2C,G). Trp and creatinine metabolite levels also changed following multiple aggression trials compared to baseline in SPF mice, displaying higher levels similar to those seen after antibiotic administration (Fig. 2D,H). DL-indole-3-lactic acid (I3LA), a metabolite of Trp, showed the same pattern but in the opposite direction. The absence of a bacterial population among the GF and ABX group led to lower levels of I3LA, and bacterial presence led to higher levels of I3LA (Fig 2E). The effect of aggression was also observed here; significantly lower levels of I3LA were observed following aggression trials compared to the baseline T0 (Fig 2F).

**Fig 2.**
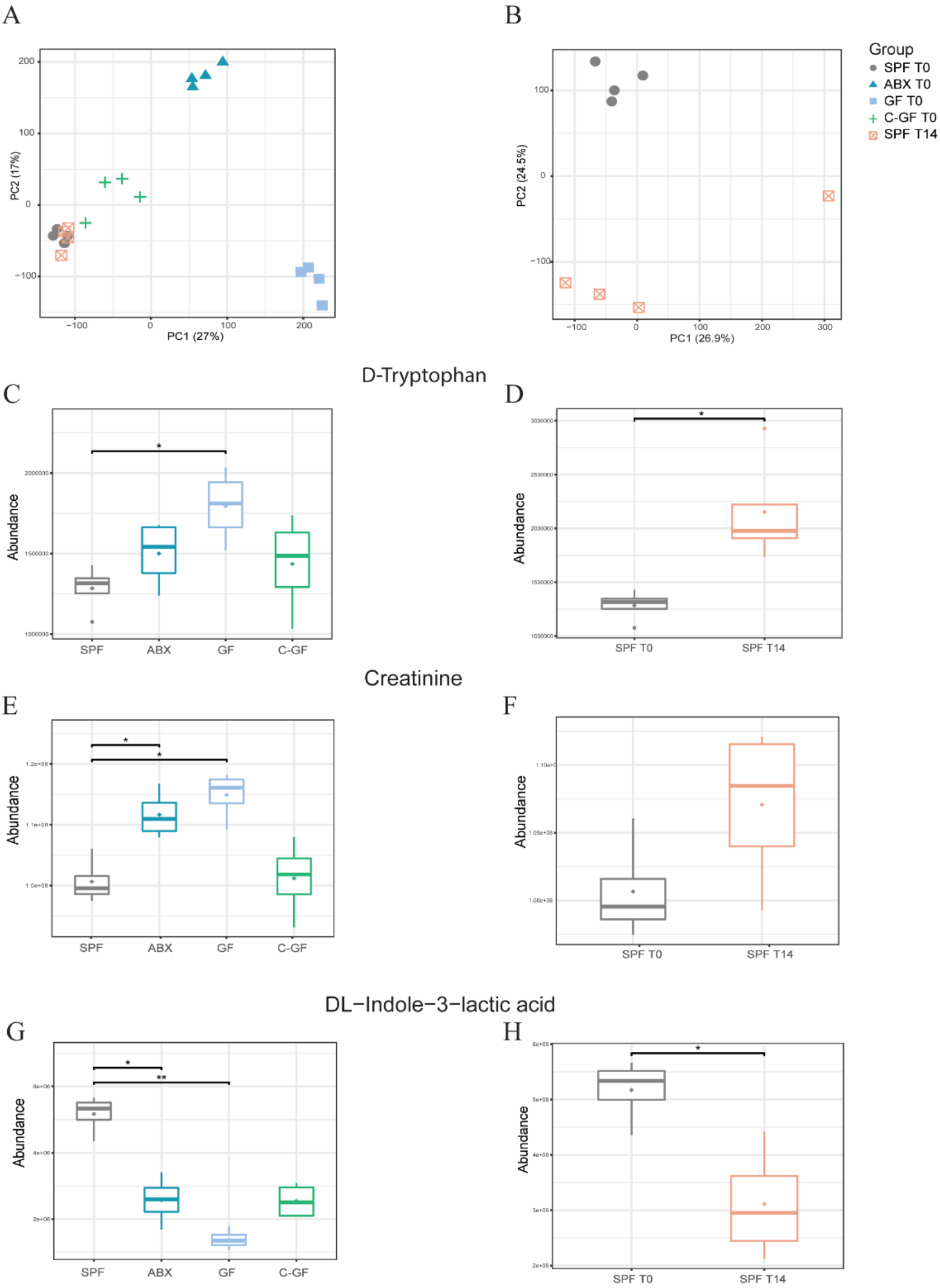
Gut microbiome and aggression induce changes in urine metabolite profiles in male mice. PCA of the overall metabolite profile in urine samples from male mice using untargeted urine metabolomics. The analysis compares two sets of samples: (A) Samples from four groups, including SPF, ABX, GF, and C-GF, taken at time point T0, before aggression tests, to assess the effect of gut microbiome perturbation on urine metabolite profiles. (B) Samples from SPF mice taken before (T0) and after (T14) aggression trials, to investigate the impact of aggression on urine metabolite profiles. Box plots illustrating the differences in (C+D) D-Tryptophan, (E+F) DL-Indole-3-lactic acid and (G+H) creatinine abundance between (C+E+G) SPF (gray), ABX (Dark blue), GF (light blue), and C-GF (green) mice and (D+F+H) SPF-T0 (gray) and SPF-T14 (orange) aggression trials. The levels are shown as normalized abundances (peak areas) (n=4).

To further our mechanistic understanding of the gut-brain-microbiome interplay, we next quantified levels of Trp, serotonin and a serotonin metabolite (5-HIAA) in the whole brain using HPLC (Fig 3A). Bacterial depletion using antibiotic treatment to the SPF group induced significant elevation in tryptophan (Trp) levels (Fig 3B) in addition to significant reduction in serotonin levels (5-HT) (Fig 3C). Furthermore, 5-HIAA and 5-HT turnover (5-HIAA/5-HT ratio), was significantly higher in the ABX group compared to the SPF group (Fig 3D and 3E respectively). These results are in line with previous studies in rats showing lower serotonin levels together with higher levels of serotonin metabolites and serotonin turnover in the brain following antibiotic treatment. In addition, Trp measured in plasma similarly showed higher levels following antibiotic treatment(*14*).

**Fig 3:**
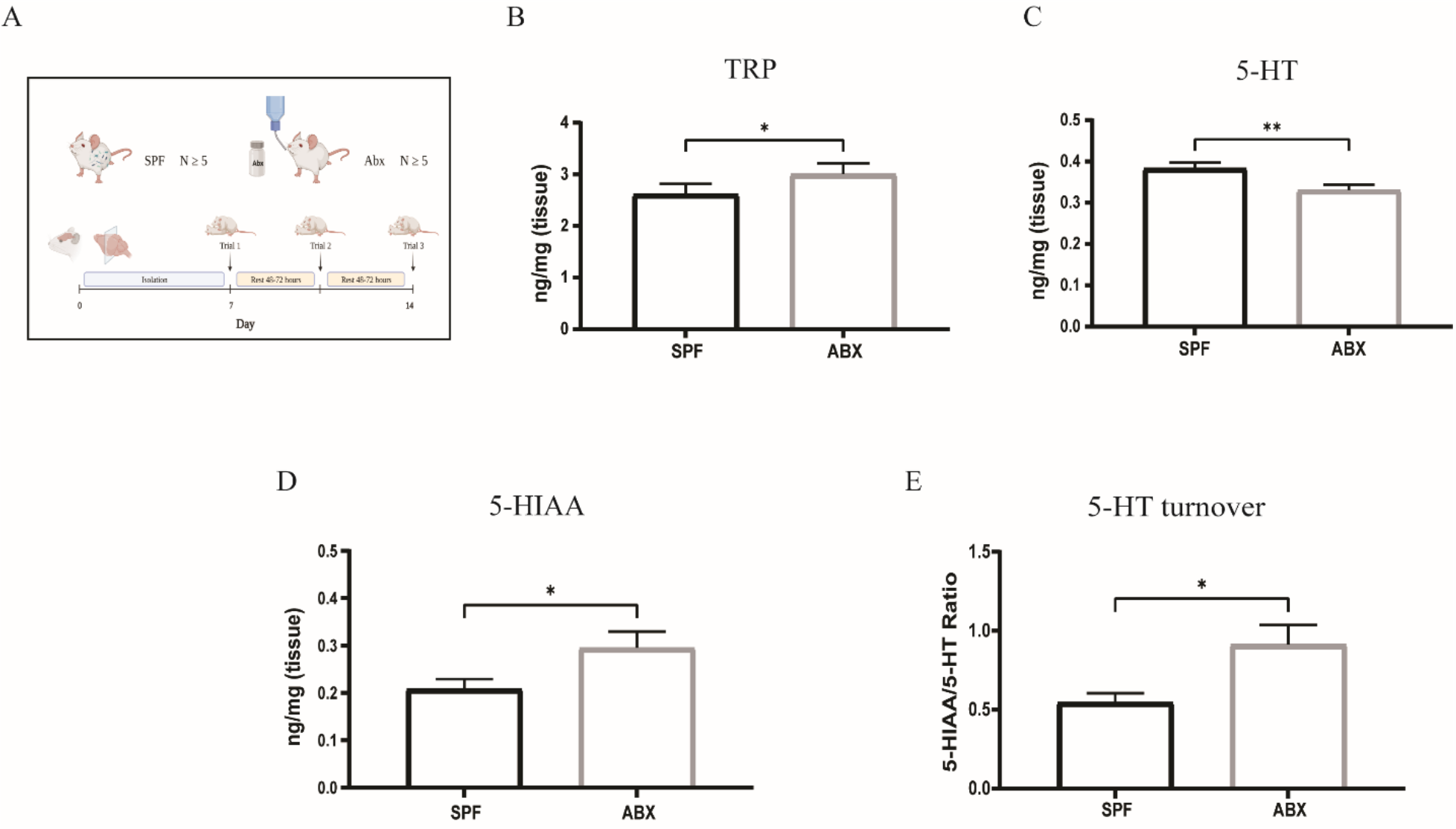
Alterations in gut bacterial composition, using antibiotic treatment, lead to changes in 5-HT, TRP, 5-HIAA and 5-HT turnover. (A) Experimental design-High pressure liquid chromatography (HPLC) performed on the whole brain of SPF and ABX treated male mice at 8 weeks of age before isolation and aggression trials. After three weeks of antibiotic treatment, we observed significant differences in the levels of (B) 5-HT, (C) TRP, (D) 5-HIAA, and (E) 5-HT turnover. (n≥5, ^*^*P*<0.05, ^**^*P*<0.01, values represent the mean +/- SEM).

A transcriptomics approach on five brain regions (prefrontal cortex, amygdala, hippocampus, hypothalamus and septum) revealed differential abundance of thousands of genes and pathways between the SPF and ABX groups (Figure 4 A-E and Table S1). We analyzed our data focusing on the 40 aggression-related genes published by Zhang-James et al.,(*6*) and key 5-HT-related genes(*15*) (Figure 4 F-J). Within this set of genes, we identified aggression-related genes that changed significantly following antibiotic treatment. Notably, the expression levels of several members of the serotonin receptor superfamily, including serotonin receptor genes 1A (htr1A), 1B (htr1B), and 2A (htr2A), were significantly altered in at least three brain regions after antibiotic treatment (Fig 4F-J). Previous studies have demonstrated that these genes play a role in aggression regulation as they encode factors that can modulate serotonergic neurotransmission. A study on silver foxes found that domesticated foxes have significantly lower density of 5-HT1A in hypothalamic membranes compared to their wild counterparts(*16*). Additionally, knockout mice lacking 5-HT1B receptors demonstrate higher aggression levels than wild-type mice(*17*) and different studies have reported an association between the HTR2A gene and aggression in both mice and humans(*18, 19*). RBFOX1, which encodes a splicing factor involved in the alternative splicing of extensive neuronal gene networks essential for brain development(*20*), was expressed at significantly higher levels in the septum following antibiotic treatment compared with the SPF group, and therefore emerges as another potential candidate for regulating aggression. Interestingly, the expression of 15 of the top 40 genes mentioned above is regulated by the protein encoded by RBFOX1(*21*). These findings provide valuable insights into the complex gut-brain interplay between the gut microbiome, gene expression, and aggression.

**Fig 4:**
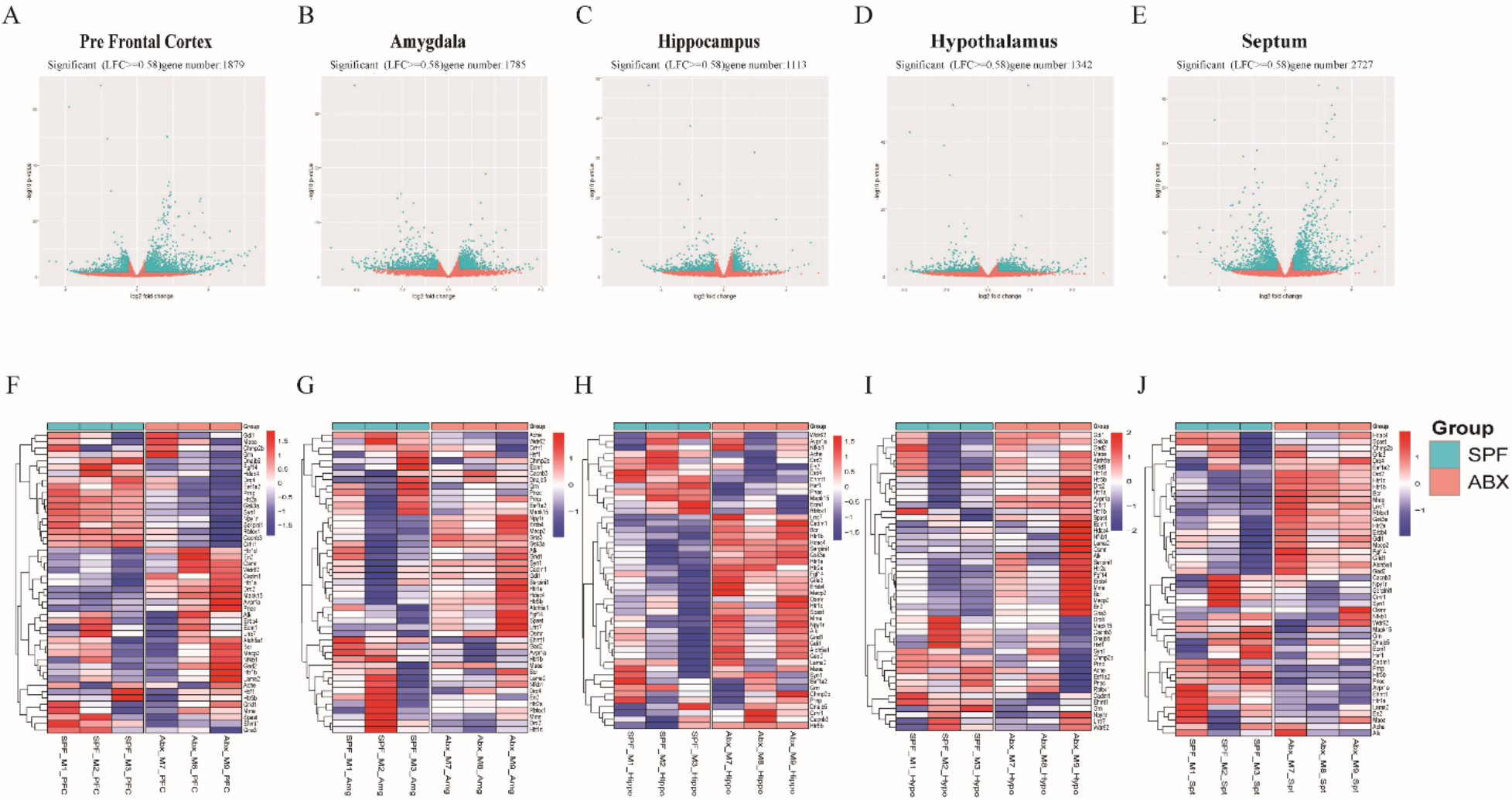
Antibiotic-induced alterations in gut bacterial composition impact gene abundance in five brain regions. RNA sequencing analysis was conducted on two groups of mice: Specific SPF (blue) and ABX (pink). The study focused on five distinct brain regions: Prefrontal cortex (A+F), Amygdala (B+G), Hippocampus (C+H), Hypothalamus (D+I), and septum (E+J). (A-E) Volcano plots displaying the differential gene expression patterns between the SPF and ABX groups in each brain region. The blue dots represent thousands of differentially expressed genes that were identified as being significantly upregulated (on the right) or downregulated (on the left) in the ABX group compared to the SPF group. The pink dots represent insignificant differentially expressed genes. (F-J) Heatmaps illustrate the relative expression levels (z-score), of 47 genes linked to aggression across the five brain regions for both SPF and ABX groups.

Following the identification of aggression-related genes that changed significantly following antibiotic treatment, the next step was to perform Gene Set Enrichment Analysis (GSEA) to identify gene sets associated with aggression (Fig 5). The analysis identified numerous enriched pathways, in all brain regions, that were significantly up or downregulated following antibiotic treatment. We found that the Rho GTPase pathway was one of the top 20 most enriched pathways in the hippocampus (Fig 5A,B, q=0.01), while the Reelin pathway showed significant enrichment in the amygdala (Fig 5C,D, q=0.003). These results align with Zhang-James *et al*.(*6*), who associated the function of Rho GTPase and Reelin pathways with aggression. Although these pathways may not have a direct relationship with aggression, they are involved in various cellular processes in the brain. Rho GTPase signaling plays a crucial role in regulating various aspects of neuronal development, growth, and survival(*22*), and the Reelin signaling pathway is crucial for nervous system development and has been linked to various neuropsychiatric disorders(*23*). In addition to these pathways, it is worth noting that serotonin-related pathways were also identified, although they did not rank among the top 50 enriched pathways. Serotonin neurotransmitter release cycle in the amygdala (q=0.03) and serotonin receptor activity in the hippocampus (q=0.04) (data not shown) were among the other significantly enriched pathways.

**Fig 5:**
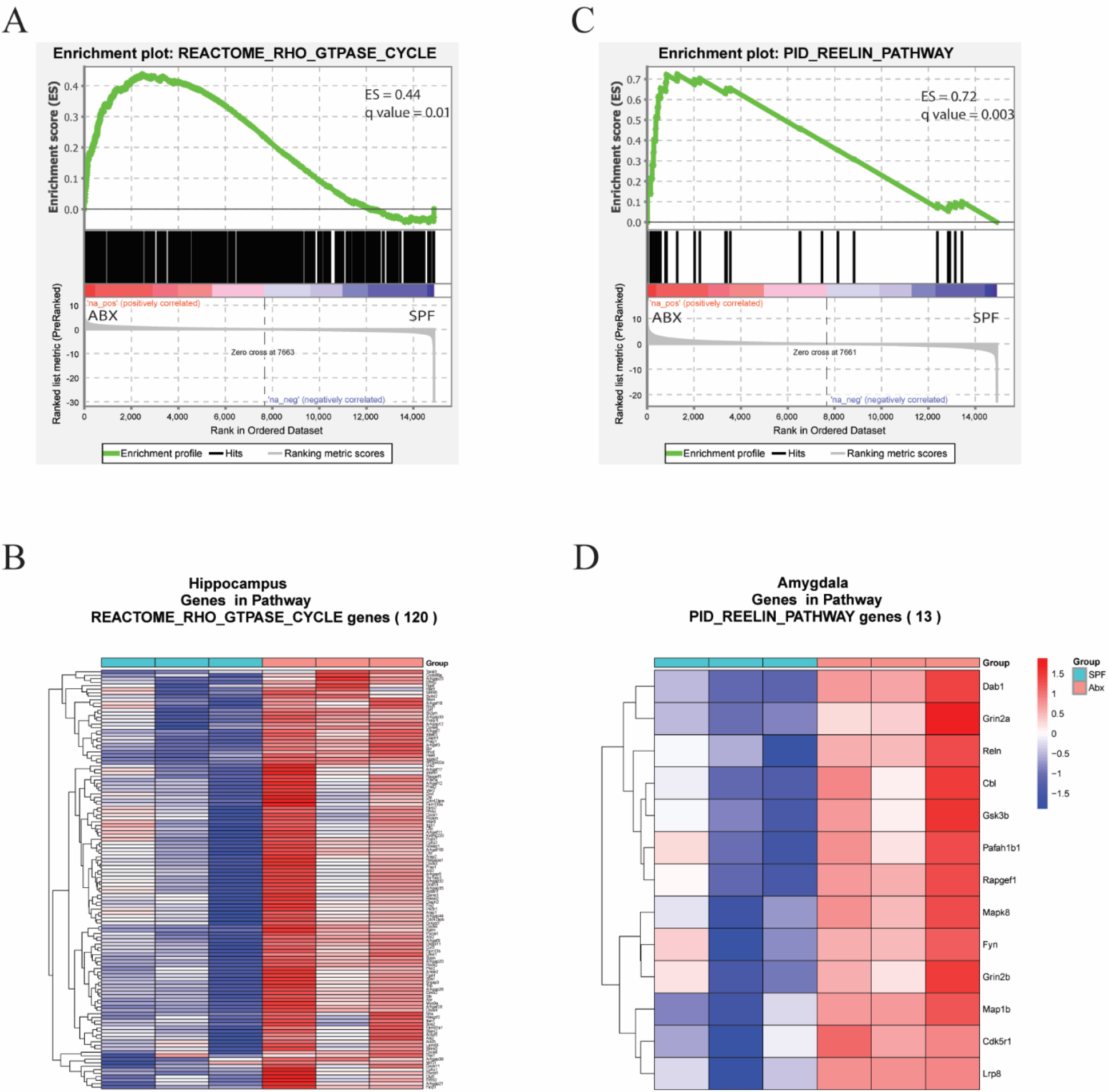
Antibiotic-induced alterations in gut bacterial composition led to changes of hundreds of pathways in five brain regions. Gene Set Enrichment Analysis, based on RNA sequencing, was employed to identify enriched pathways. (A+C) enrichment plot of Rho GTPase cycle in the hippocampus and Reelin pathway in the amygdala, respectively, illustrate the differentially expressed genes (DEGs) identified between the SPF and ABX groups. The top portion of plots show the enrichment scores for each gene, and the bottom portion shows the ranked genes. (B+D) Heatmap displays the core enrichment genes’ relative expression (z-scores) identified for each pathway, Rho GTPase cycle and Reelin pathway.

## Early-life antibiotic use increased aggression in humanized mice

Next, we aimed to translate our findings into a clinical context, exploring how antibiotic use in early life affects aggression. Towards this end, we performed fecal microbiome transplant (FMT) from feces of one-month-old infants who were or were not exposed to antibiotics in the first 48 hours of life, into five-week-old GF mice. Impressively, behavioral assays in mature mice confirmed our above findings: antibiotic-altered microbial communities, here of infants, led to increased aggression, even when antibiotic use has ended and the microbiome has had time to recover (samples were collected 1 month after antibiotic exposure) (Figure 6).

**Fig 6:**
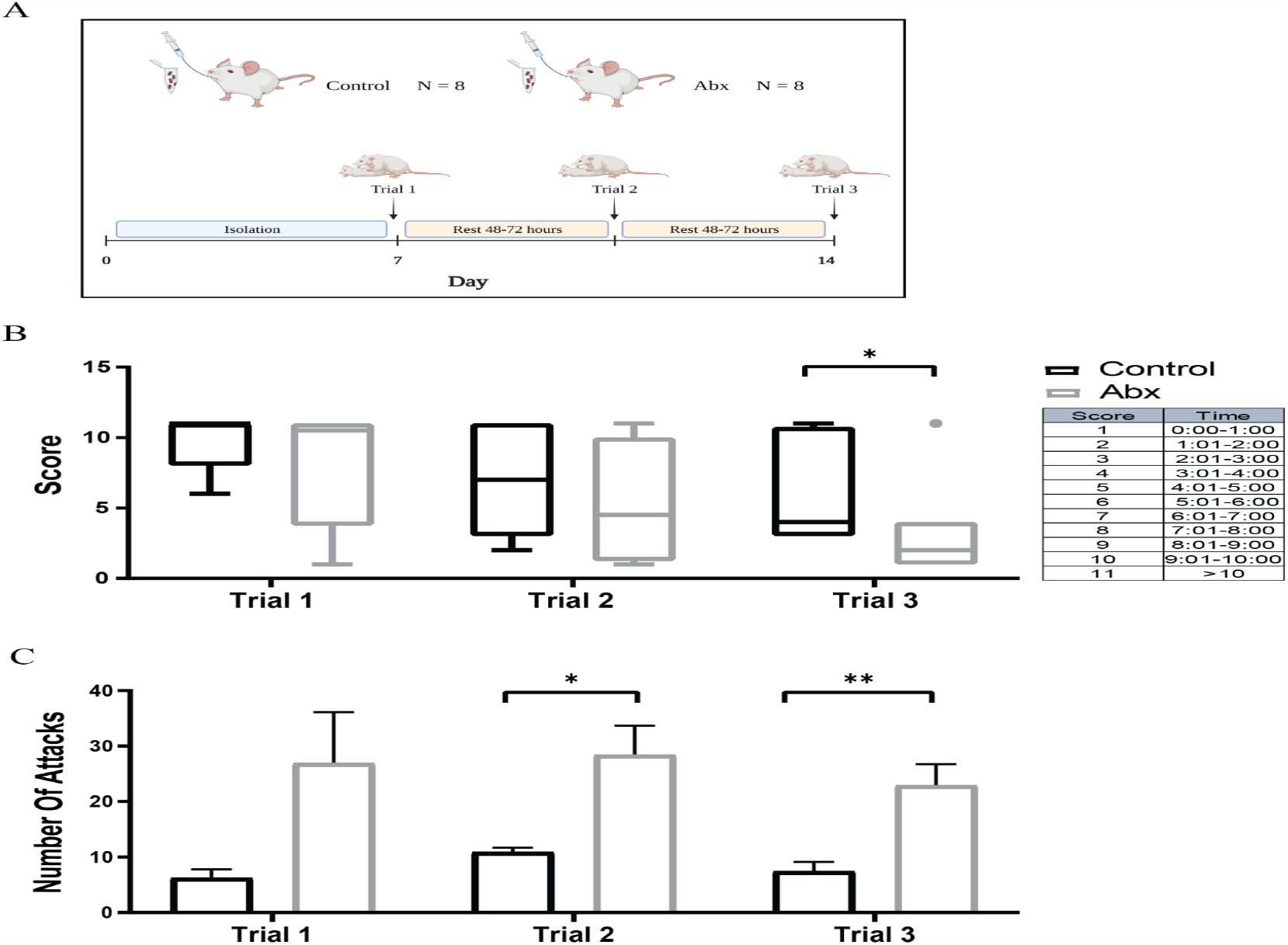
Fecal Microbiome Transplantation (FMT) from antibiotic-treated babies, collected 1 month after exposure, to GF mice induces increased aggression in mice. (A) Experimental design – FMT from antibiotic-treated (grey) and control (black) infants to five-week-old GF mice. At the age of 8 weeks, aggression was examined using the resident-intruder test as described above. (B) Attack latency, the time between the first introduction of the intruder until the first attack by the resident: every minute get a score in order to achieve a discrete and non-continuous variable, and (C) number of attacks, the overall number of attacks in each trial (n≥9, ^*^*P*<0.05, ^**^*P*<0.01, number of attacks represents the mean +/- SEM).

## Discussion

The present study provides insights into the role of the gut microbiome in modulating aggression in a murine model and also humanized mice, supporting the involvement of the gut-brain axis in regulating social behaviors. This study confirms the significant impact of microbiome on aggression, consistent with previous studies(*23*). However, our findings not only demonstrate the impact of gut microbiome on aggression but also reveal its influence on multiple factors and mechanisms that regulate this behavior. To gain a mechanistic understanding of the gut-brain-microbiome crosstalk, we first explored urine metabolite profiles using untargeted metabolomics. Our findings revealed distinct shifts in metabolite profiles following bacterial perturbation and aggression trials, including in tryptophan, creatinine, and indole-3-lactic acid. Furthermore, our analysis of serotonin levels in the whole brain, supported by findings by Hoban et al.(*14*), demonstrated lower serotonin levels and higher levels of tryptophan, serotonin metabolite, and serotonin turnover following antibiotic treatment. Moreover, gene expression analysis showed alterations in serotonin receptor genes across different brain regions following antibiotic treatment, supporting the involvement of serotonin signaling pathways in mediating the gut microbiome’s effects on aggression. Notably, the study revealed the clinical relevance of early-life antibiotic use, as FMT from infants exposed to neonatal antibiotics led to increased aggression in humanized mice. These findings highlight the complex interplay between the gut microbiome and aggression, providing valuable insights into the underlying biological processes involved. However, further research is needed to fully understand the complex interactions between the gut microbiome, gene expression, and aggression, and how these findings can be translated into clinical applications.

## Supplementary Text

### Materials and Methods

#### Mice

Swiss Webster mice were obtained from Taconic Farms Inc. (Germantown, NY, USA) and maintained at the animal facility of the Azrieli Faculty of Medicine. Mice were maintained in standard 12h:12h light:dark housing conditions; specific pathogen-free (SPF) mice and antibiotic-treated SPF mice (ABX) were housed in the SPF animal house of the Azrieli Faculty of Medicine, Bar-Ilan University (BIU); conventionalized germ-free mice (C-GF) and humanized mice were held in the conventional room of the animal house; and the GF group was housed in regular cages inside sterile isolators under the same growth conditions but with autoclaved food and water. Both male and female mice were used in this study, which was performed using protocols approved by the Bar Ilan University Animal Studies Committee.

#### Antibiotic-treated mice

For the antibiotic-treated SPF group, mice at the age of 5 weeks were treated with a combination of ciprofloxacin (0.04 gl^-1^) (Sigma-Aldrich Corporation, St. Louis, MO, USA), metronidazole (0.2 gl^-1^) (Santa Cruz Biotechnology, Santa Cruz, CA) and vancomycin Hudrochlorid (0.1 gl^-1^) (Gold Biotechnology, St. Louis, MO, USA) in drinking water, refreshed twice a week for 3 weeks before and during the experiment.

#### Re-conventionalized mice

At five weeks of age, GF mice were colonized by oral transplantation with stool samples collected from SPF mice of the same age. Stool samples were suspended in sterile phosphate-buffered saline (PBS) (1 fecal pellet/1 ml of PBS) and dissolved by vortex for 1 minute. A total of 150 μl of the fecal suspension was administered by oral gavage to GF mice. The process was performed once, immediately after mice were removed from the sterile isolator, followed by housing in the conventional animal facility under the same conditions as SPF and GF mice.

### Humanized mice

#### Fecal sample donor

Fecal samples were obtained from subjects from a clinical study conducted at the Turku University Hospital in Turku, Finland(*24*). Altogether, 13 infants exposed to antibiotic therapy with intravenous benzylpenicillin and gentamicin during the first 48 hours of life and 20 non-exposed infants were selected (as controls) based on sample availability. The samples (taken when infants were one month old) were maintained at -80°C and shipped on dry ice to the Azrieli Faculty of Medicine, BIU. Enrollment criteria are presented by Uzan-Yulzari et al.(*24*), and the study was approved by the Finnish Institute for Health and Welfare, a national expert agency under the jurisdiction of the Finnish Ministry of Social Affairs and Health(*24*).

#### Fecal transplantation

Fecal samples from antibiotic-treated and control infants were transplanted to GF male mice by oral gavage at 5 weeks of age. Each sample was suspended in 800μl of sterile phosphate buffered saline (PBS) and dissolved by vortex for 1-minute. A total of 200 μl of the fecal suspension was administered by oral gavage to one to three GF male mice. The process took place once, immediately after the mice were taken out of the isolator, followed by housing in the conventional animal facility under the same conditions as SPF and GF mice.

#### Resident intruder test

Aggression was measured using the resident-intruder paradigm test(*13*) among all four groups of mice: SPF, GF, colonized GF, and antibiotic-treated SPF, between 8-12 weeks of age. Mice were tested three times at 2-4 day intervals. All trials were carried out during the light phase of the light-dark cycle between 12-3 pm. Each male was housed with a companion female for one week before the start of the experiments; females were removed from the residential cage one hour before the trial. Each resident was tested in his home cage against a group-housed intruder for 10 minutes. After completion of the trial, the intruder male was removed from the cage and the companion female was reunited with the resident male until the next trial. Aggression was measured using two parameters: attack latency and number of attacks. Attack latency was scored as the time to first aggressive attack; mice that attacked during the first minute were scored 1, during the second minute were scored 2 and so on. Initial statistical comparisons for mouse aggression tests were performed by Kruskal-Wallis, with uncorrected Dunn post-hocs (SPF, GF, C-GF, ABX) and subsequent two-group comparisons (SPF-ABX, humanized ABX-humanized control) were made with one-tailed Mann Whitney U test, using Prism 9.5.0 (GraphPad Software, San Diego, CA, USA).

### Untargeted Metabolomics of Mouse Urine

#### Sample preparation

Urine samples were collected from male mice between 8-9 weeks of age directly into 1.5ml eppendorf tubes using the bladder massage method(*14*). Samples were maintained at -80°C and shipped on dry ice to Afekta Technologies Ltd., Finland. Upon arrival, mouse urine samples were thawed in an ice-water bath, vortexed (10 s), and for dilution, aliquots were transferred into filter plate (Captiva ND filter plate 0.2 μm) and Class 1 ultrapure water was added and mixed in a ratio of 300 μl per 100 μl of sample. For metabolite extraction, cold acetonitrile was added in a ratio of 400 μl per 100 μl per of urine sample and mixed. The samples were then centrifuged for 5 min at 700×g at 4 °C and kept at 4 °C until analysis. The pooled quality control (QC) sample was prepared by collecting 50 μl from each supernatant and combining the material in one tube.

#### LC–MS analysis

The samples were analyzed by liquid chromatography–mass spectrometry (LC-MS), consisting of a 1290 Infinity II UHPLC (Agilent Technologies, Santa Clara, USA) coupled with a high-resolution QTOF mass spectrometer (Agilent 6546 with Jet Stream ion source, Agilent Technologies). The analytical method has been described previously(*25, 26*). In brief, a Zorbax Eclipse XDB-C18 column (2.1 × 100 mm, 1.8 μm; Agilent Technologies) was used for the reversed-phase (RP) separation and an Aqcuity UPLC BEH amide column (Waters) for the HILIC separation. After each chromatographic run, the ionization was carried out using jet stream electrospray ionization (ESI) in the positive and negative mode, yielding four data files per sample. The collision energies for the MS/MS analysis were selected as 10, 20, and 40 V, for compatibility with spectral databases.

#### Data preprocessing

Peak detection and alignment were performed in MS-DIAL ver. 4.90(*27*). For the peak collection, m/z values between 50 and 1500 and all retention times were considered. The amplitude of minimum peak height was set at 5000. The peaks were detected using the linear weighted moving average algorithm. For the alignment of the peaks across samples, the retention time tolerance was 0.2 min and the m/z tolerance was 0.015 Da. Solvent background was removed using solvent blank samples under the condition that to be kept for further data analysis, the maximum signal abundance across the samples had to be at least five times that of the average in the solvent blank samples.

After the peak picking, 86,194 detected molecular features were included in the data preprocessing and clean-up step. Low-quality features were flagged and discarded from statistical analyses. Molecular features were only considered high-quality if they met all the following quality criteria: low number of missing values, present in more than 70% of the QC samples, present in at least 60% of samples in at least one study group, RSD^*^ below 20%, D-ratio^*^ below 10%. In addition, if either RSD^*^ or D-ratio^*^ was above the threshold, the features were still considered high-quality if their classic RSD, RSD^*^ and basic D-ratio were all below 10%. The signals were normalized for signal drift and batch effect. After the preprocessing and data clean-up, 62837 molecular features were considered high-quality and included in the FDR correction calculations. The high number of molecular features before data clean-up is due to the high sensitivity of the instrument, collecting several signals from each actual metabolite, but also from the solvent background and detector noise.

## Data analysis

For statistical analyses of urine sample metabolite profiles, we used feature-wise Welch’s t-tests. Fold changes (relative change or difference between two groups as a ratio of the group means (the first divided by the latter)) and Cohen’s D (a measure of effect size as a difference in standard deviations) values were computed to measure the effect size in each analysis. P-values of high-quality features were adjusted for multiple testing with FDR correction. All analyses were conducted with R version 4.1.2.

### Compound identification

The chromatographic and mass spectrometric characteristics (retention time, exact mass, and MS/MS spectra) of the significantly differential molecular features were compared with entries in an in-house standard library and publicly available databases, such as METLIN and HMDB, as well as with published literature. The annotation of each metabolite and the level of identification was given based on the recommendations published by the Chemical Analysis Working Group (CAWG) Metabolomics Standards Initiative (MSI)(*28*).

### High performance liquid chromatography of whole brains (HPLC)

To determine levels of serotonin, tryptophan, and associated metabolite 5-HIAA, we performed high performance liquid chromatography (HPLC) analysis on whole brains from male mice 8-9 weeks of age that did not undergo behavioral assays (sacrificed by rapid decapitation) and prepared as described in(*29*).

Tissue levels of TRP, 5-HT and its metabolite 5-HIAA (Cat Nrs. T0254, 14927, 55697, respectively Merck KGaA, Darmstadt, Germany) were analyzed using uHPLC with fluorometric detection.

### Statistical analysis

Amounts of 5-HT, TRP and 5-HIAA were normalized to the wet tissue weight for statistical analysis and calculation of substance levels was based on external standard values. Statistical comparisons of 5-HT, TRP, 5-HIAA and 5-HT turnover were performed by one-tailed Mann Whitney U test, using Prism 9.5.0 (GraphPad Software, San Diego, CA, USA). Graphs are presented as mean ± SEM, asterisks indicate significance ^(*^p<0.05,^**^p<0.01, ^***^p<0.0001).

### RNA sequencing (transcriptomics)

Brain punches were taken from sacrificed SPF and antibiotic-treated SPF mice that did not undergo behavioral testing between 8-9 weeks of age. Mice were sacrificed by live rapid decapitation and brains were removed. Five regions of interest were dissected and frozen on dry ice: the hippocampus, prefrontal cortex and septum were isolated using a 13 gauge needle, the amygdala using a 16 gauge needle, and the hypothalamus using a 14 gauge needle. Samples were kept in - 80°C until processing.

Total RNA was extracted using the RNAeasy Mini Kit according to the manufacturer’s instructions (QIAGEN, Manchester, UK). Integrity of the isolated RNA was tested using the Agilent RNA Pico Kit and Bioanalyzer at the Genome Technology Center at the Azrieli Faculty of Medicine, BIU. Total RNA was taken for mRNA isolation using a NEBNext Poly(A) mRNA Magnetic Isolation Module (New England Biolabs, Inc., Ipswich, MA, USA)) and libraries were prepared using the NEBNext Ultra II RNA Library Prep Kit for Illumina (New England Biolabs, Inc., Ipswich, MA, USA)). Quantification of the library was performed using dsDNA HS Assay Kit and Qubit 2.0 (Molecular Probes, Life Technologies). 4nM of the library was denatured in 0.2M NaOH for 5 min at room temperature; 1.45pM was loaded onto the Flow Cell with 1% PhiX library control.

Libraries were sequenced on an Illumina NextSeq550 instrument, 75 cycles single-read sequencing.

### RNA data analysis

For RNA sequencing data analysis, reads were aligned to the *Mus musculus* reference genome GRCm39 using STAR (version 020201)(*30*), and quantification of reads was performed using htseq-count (version 0.12.4)(*31*) and a list of genes (Ensembl gtf file)(*32*). Differential gene expression analysis was then performed using DESeq2 (version 1.30.1)(*33*). Significant differentially expressed genes were selected using threshold values of p-value smaller than 0.05 and log_2_fold change greater or equal to 0.58. A volcano plot was rendered using ggplot2 (version 3.3.2) and heatmaps were generated using pheatmap (version 1.0.12)(*34*). For pathway enrichment analysis, Gene Set Enrichment Analysis (GSEA version 4.0.3)(*35*) was used for all the genes ranked (-log_10_(pvalue)/sign(log_2_FoldChange)), converted to human genes, using three datasets: hallmark, Curated Canonical Pathways and GO gene sets. GSEA results(*q*-value<=0.05) were categorized and plotted (using ggplot2 version 3.3.2).

Tables S1

https://docs.google.com/spreadsheets/d/1vnhZQryRNtzB2SpMTJTfyAI-MacHD2bB/edit?usp=sharing&ouid=118053953844866984682&rtpof=true&sd=true

## Notes

### Competing Interest Statement

The authors have declared no competing interest.

